# Loss of the polarity protein Par3 promotes dendritic spine neoteny and enhances learning and memory

**DOI:** 10.1101/2023.08.30.555530

**Authors:** Mikayla M. Voglewede, Elif Naz Ozsen, Noah Ivak, Matteo Bernabucci, Miao Sun, Zhiping P. Pang, Huaye Zhang

## Abstract

The Par3 polarity protein is critical for subcellular compartmentalization in different developmental processes. Variants of *PARD3*, which encodes PAR3, are associated with intelligence and neurodevelopmental disorders. However, the role of Par3 in glutamatergic synapse formation and cognitive functions *in vivo* remains unknown. Here, we show that forebrain conditional knockout of Par3 leads to an increase in long, thin dendritic spines without significantly impacting mushroom spines *in vivo*. In addition, we observed a decrease in the amplitude of miniature excitatory postsynaptic currents. Surprisingly, loss of Par3 *in vivo* enhances hippocampal-dependent spatial learning. Phosphoproteomic analysis revealed proteins regulating cytoskeletal dynamics are significantly dysregulated downstream of Par3. Mechanistically, we found Par3 deletion causes increased activation of the Rac1 pathway. Together, our data reveal an unexpected role for Par3 as a molecular gatekeeper in regulating the pool of immature dendritic spines, a rate-limiting step of learning and memory, through modulating Rac1 activation *in vivo*.

## Introduction

Dendritic spines are small, highly polarized protrusions on neurons serving as sites for most excitatory postsynaptic input. Plasticity of dendritic spines is necessary for learning^1–7^, while stable dendritic spines are thought to encode long-term memories^8^. In the dendritic spine head, the protein rich postsynaptic density (PSD) serves as the incoming signaling center opposing the presynaptic terminal. The PSD contains neurotransmitter receptors and signaling molecules crucial for synaptic transmission^9^, making it a distinct zone compared to the cytoskeleton rich dendritic spine neck and dendrite shaft. Both actin and microtubules provide structure and contribute to the function of the compartmentalized domains within the dendritic spine^10,11^. However, the molecular mechanisms governing the cytoskeletal dynamics in the morphogenesis of these polarized spine structures remain incomplete.

The partitioning defective (Par) polarity complex, including Par3, a scaffolding molecule, Par6, an adaptor molecule, and atypical protein kinase C (aPKC), is responsible for establishing cell polarity in various cellular contexts, such as apical-basal compartmentalization and planar cell polarity^12–14^. In primary hippocampal neurons, knockdown of Par3 or Par6 causes a drastic shift to immature dendritic spines with a loss of presynaptic partners^15,16^. Par3 directly interacts with TIAM1, a guanine nucleotide exchange factor for Rac GTPases, and restricts Rac activity to the dendritic spines^15^. Together, Par3 and TIAM1 are recruited to the synapses by and directly interact with the adhesion GPCR, brain-specific angiogenesis inhibitor 1 (BAI1)^17^. In addition, double conditional knockout of both aPKC isozymes, PKCι/λ and PKCζ, leads to impaired learning and memory^18^.

Interestingly, a copy number variant of *PARD3*, the gene that encodes PAR3, is associated with autism spectrum disorder (ASD)^19^. Various single nucleotide polymorphisms (SNPs) of *PARD3* are associated with schizophrenia^20^, intelligence^21^, education attainment^22^, and high math ability^22^. Similarly, *PARD3B,* a paralog of *PARD3*, is associated with ASD^23–27^. This suggests that PAR3 may play a key role in higher cognition and behavior. However, the *in vivo* role for Par3 in synaptic function and cognition remains completely unknown. Thus, we created a conditional knockout mouse model to postnatally knockout *Pard3* in forebrain pyramidal neurons to investigate the role of Par3 in dendritic spine morphogenesis and learning and memory *in vivo*. Here, we show the loss of Par3 *in vivo* creates excess immature dendritic spines in the CA1 of the hippocampus and leads to a significant decrease in the amplitude of miniature excitatory postsynaptic currents (mEPSCs). Interestingly, the loss of Par3 enhances spatial learning and memory and dysregulates the Rac1 pathway and cytoskeletal signaling. These data reveal a surprising role for Par3 in regulating a rate-limiting step of learning and memory through limiting the pool of immature dendritic spines.

## Results

### Generating the Pard3 conditional knockout model

To examine the effects of Par3 on dendritic spine morphogenesis and learning and memory *in vivo*, we established a *Pard3* conditional knockout mouse model by crossing *Pard3^<tm1a(KOMP)Wtsi>^*/H mice with *CAG-Flpo* mice. The resulting *Pard3^f/f^* line (Tm1c) was crossed with *CaMKIIα-Cre* line through which Cre excises exons 8 and 9 of *Pard3* (Figure S1A). Since Par3 plays a role in establishing neuronal polarity^28^ and to avoid early developmental defects, *CaMKIIα-Cre* was chosen to knockout *Pard3* two to three weeks postnatally. The presence of floxed *Pard3* gene was validated via PCR (Figure S1B). Cre expression was confirmed by crossing the *Pard3^f/f^* and *Pard3^f/f^:CaMKIIα-Cre* mice with B6.129X1-*Gt(ROSA)26Sor^tm1(EYFP)Cos^*/J, in which enhanced yellow fluorescent protein (EYFP) is expressed in *Cre*-expressing tissue. CaMKIIα-Cre is highly expressed in the hippocampus, specifically the CA1, with sparse cortical expression (Figure S1C), supporting previous publications using *CaMKIIα-Cre*^29,30^. In primary neurons derived from E17.5 *Pard3^f/f^* infected with AAV-Cre-GFP-hSyn, the Par3 protein was ablated (Figure S1D). Additionally, conditional knockout of Par3 did not alter body weight (Figure S1E). This novel mouse model provides a unique opportunity to investigate the role of Par3 in dendritic spines and cognition *in vivo*.

### Loss of Par3 promotes neoteny of dendritic spines

*In vitro* shRNA-mediated knockdown of Par3 in primary hippocampal neurons results in an increase in immature, filopodia-like spines and a decrease in mature dendritic spines^15^. By contrast, the loss of Par3 *in vivo* increases the density of dendritic spines in the stratum radiatum layer of hippocampal CA1 pyramidal neurons in both male and female mice (Figure 1A-B and S2A-B, males: p= 0.0226; females: p= 0.0246). Interestingly, female mice exhibit higher density of dendritic spines compared to age-matched males, which is consistent with previous reports showing higher dendritic spine density in proestrus females compared to males^31^, even though estrous cycle was not determined in our experiments. Overall, our results demonstrate that loss of Par3 increases dendritic spine density regardless of sex.

**Figure 1:**
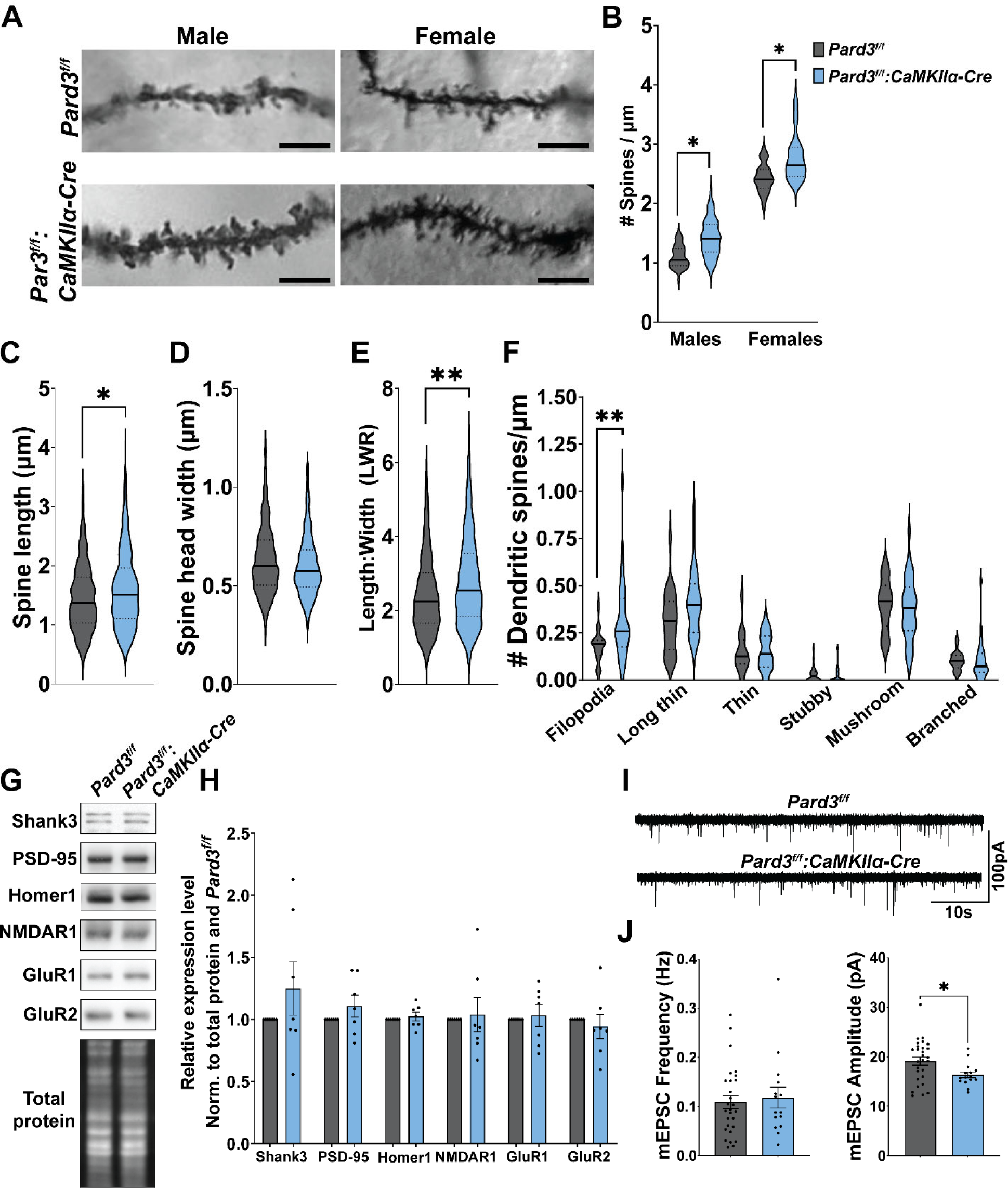
Loss of Par3 increases density of longer, thinner dendritic spines and alters synaptic function. See also figure S2. (A) Representative images of *Pard3^f/f^*(top) and *Pard3^f/f^:CaMKIIα-Cre* (bottom) dendritic spines. Scale bar = 5 µm. (B) Quantification of dendritic spine density. N = 38 total dendrites from 3 *Pard3^f/f^* males, 58 total dendrites from 5 *Pard3^f/f^:CaMKIIα-Cre,* males, 30 total dendrites from 3 *Pard3^f/f^*females, and 29 total dendrites from 3 *Pard3^f/f^:CaMKIIα-Cre* females. (C-F) Quantification of (C) spine length, (D) spine head width, (E) length to width ratio. N = 1164 total spines from 3 *Pard3^f/f^* mice and 1407 total spines from 5 *Pard3^f/f^:CaMKIIα-Cre* mice. (F) Quantification of morphology classification. N = 30 total dendrites from 3 *Pard3^f/f^* mice and 36 total dendrites from 5 *Pard3^f/f^:CaMKIIα-Cre* mice. In B-F, data representation includes violin plots with thick lines as medians and dotted lines as quartiles. Data are analyzed via mixed model analysis. *p≤0.05, **p≤0.01. (G) Representative images for proteins detected from PSD-enriched fractions derived from *Pard3^f/f^* or *Pard3^f/f^:CaMKIIα-Cre* hippocampal tissue. (H) Quantification of relative levels of proteins in PSD-enriched fractions normalized to total protein and to *Pard3^f/f^*including SH3 and multiple ankyrin repeat domains 3 (Shank3), postsynaptic density protein 95 (PSD-95), homer scaffold protein 1 (Homer1), N-methyl-D-aspartate (NMDA) receptor subunit 1 (NMDAR1), glutamate ionotropic receptor AMPA type subunit 1 (GluR1) and 2 (GluR2). N = 7 *Pard3^f/f^* mice and 7 *Pard3^f/f^:CaMKIIα-Cre* mice. (I) mEPSC traces and (J) quantification of mEPSC frequency and amplitude. N = 28 *Pard3^f/f^* cells and 15 *Pard3^f/f^:CaMKIIα-Cre* cells. In G-J all data are presented as individual mice or cells as points and bar graphs as mean ±SEM, analyzed with unpaired T-test. *p≤0.05

We then analyzed dendritic spine morphology in *Pard3^f/f^*and *Pard3^f/f^:CaMKIIα-Cre* hippocampi. Dendritic spine length (Figure 1C) and spine head width (Figure 1D) were used to calculate length to width ratio (LWR, Figure 1E) in *Pard3^f/f^* and *Pard3^f/f^:CaMKIIα-Cre* males. The length, width, and LWR were used to classify spines into morphological categories. The loss of Par3 results in longer spines (Figure 1C, p=0.0149) with an increased LWR (Figure 1E, p=0.0033) but no significant change in spine head width (Figure 1D). The increased LWR of *Pard3^f/f^:CaMKIIα-Cre* dendritic spines is characteristic of a shift to long, thin spines more commonly found in early development. Surprisingly, the density of mature dendritic spines did not change. However, there was an increase in filopodia spines (Figure 1F, p=0.0027) and a trend toward increased long thin dendritic spines (Figure 1F, p=0.1075). Together, this indicates the loss of Par3 results in a shift towards longer, thinner dendritic spines and increased dendritic spine density commonly found in early development prior to synaptic refinement and maturation.

We next aimed to determine how the extra pool of long, thin dendritic spines alter synaptic protein expression and function. We isolated PSD-enriched fractions from hippocampal tissue using centrifugation (Figure S2C). Although the complete elimination of the presynaptic membrane does not occur, PSD-related proteins are enriched providing a cleaner investigation of synaptic changes (Figure S2D). In the PSD-enriched fraction, the loss of Par3 does not result in altered synaptic levels of proteins including SH3 and multiple ankyrin repeat domains 3 (Shank3), post synaptic density protein 95 (PSD-95), homer scaffold protein 1 (Homer1), N-methyl-D-aspartate receptor subunit 1 (NMDAR1), AMPA-type glutamate receptor subunit 1 (GluR1), or subunit 2 (GluR2) (Figure 1G). Drawing a definite conclusion is challenging because the PSD-enriched fraction may not encompass the surplus of filopodia and long thin dendritic spines, which normally have minimal or no PSDs. Thus, the similar synaptic protein expression could be due to similar density of mature dendritic spines (Figure 1F), or it could be due to similar amounts of synaptic proteins spread across the increased dendritic spine density.

Next, our objective was to investigate potential changes in synaptic transmission in the *Pard3^f/f^:CaMKIIα-Cre* mice. Basal excitatory synaptic transmission was measured by recording miniature excitatory postsynaptic currents (mEPSCs) in the CA1 of the hippocampus of 5- to 7- week-old mice. Interestingly, we found a significant decrease in the amplitude of mEPSCs with no significant changes in mEPSC frequency in the *Pard3^f/f^:CaMKIIα-Cre* hippocampus (Figure 1I- J, amplitude: p=0.0271), which suggests the loss of Par3 in the hippocampus decreases synaptic strength. This decrease in mEPSC amplitude is consistent with the observation that *Pard3^f/f^:CaMKIIα-Cre* neurons have increased density of filopodia and long thin spines. Since these immature synaptic features are common in early postnatal development, our data suggest the loss of Par3 promotes dendritic spine neoteny, leading to the retention of immature synaptic features in early adulthood.

### Loss of Par3 enhances learning and memory

We next investigated whether the increase in immature dendritic spines alters cognitive function, specifically hippocampal-dependent spatial learning via the Morris Water Maze (MWM). In the MWM mice underwent 5 trials per day on 4 consecutive training days to learn the location of a hidden platform in opaque water using spatial cues. 24 hours after the last training day, memory was tested during the probe test. In the probe test the platform was removed and mice swam for 60 seconds. The time spent in each quadrant and the distance from platform location were used as indicators of spatial memory retention (Figure 2A). Surprisingly on training day 2 of the MWM, *Pard3^f/f^:CaMKIIα-Cre* mice reached the hidden platform ∼39% more quickly than *Pard3^f/f^* mice (Figure 2B, day 2: p=0.0191), suggesting either they retained a stronger memory of the platform location from training day 1 and/or their learning capacity was enhanced. This was not due to increased exposure leading to enhanced learning and memory, as *Pard3^f/f^* and *Pard3^f/f^:CaMKIIα- Cre* exhibited similar swimming velocities during all training days (Figure S3C). Throughout the training period, there were no significant sex differences (Figure S3A-B). During the probe test, both *Pard3^f/f^* and *Pard3^f/f^:CaMKIIα-Cre* learned the location of the platform as indicated by spending similar amounts of time in and cumulative distance from the target quadrant where the platform was previously located (Figure 2C and E, S3D-E). Interestingly, we observed a significant behavioral difference during the training sessions. After successfully reaching the platform, some mice attempted jumping from the platform towards the maze wall. Surprisingly, 54.5% of the *Pard3^f/f^:CaMKIIα-Cre* mice jumped from the platform, nearly all of which jumped during 3 or more trials. Only 21% of *Pard3^f/f^* mice jumped from the platform, nearly all of which jumped at the end of only a single trial. Overall, *Pard3^f/f^:CaMKIIα-Cre* mice attempted jumping at the end of more trials than *Pard3^f/f^* (Figure 2D, p=0.0446). This is intriguing as repeated jumping behavior has been observed in ASD mouse models such as the Shank2 knockout mice^32^.

**Figure 2:**
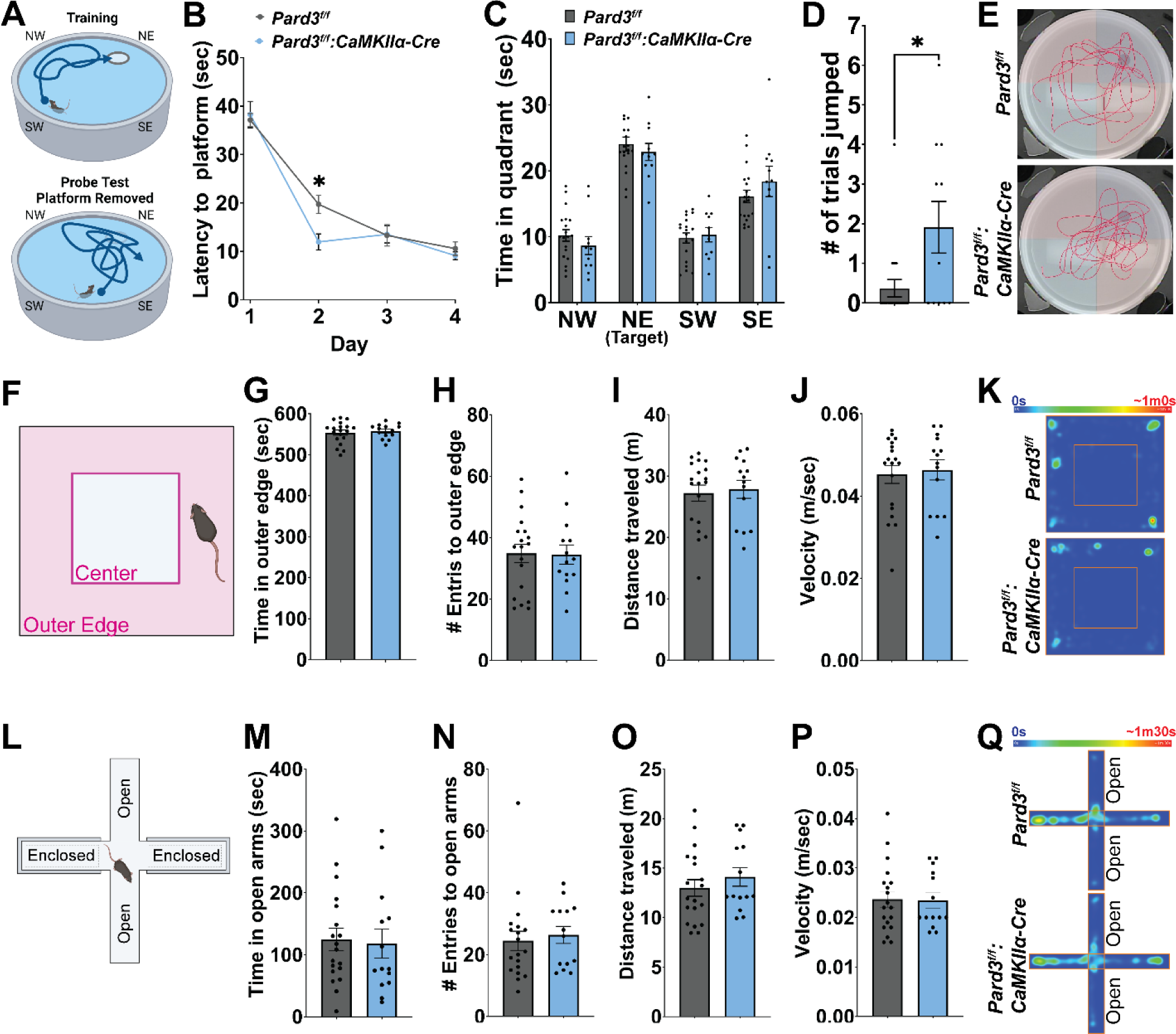
Loss of Par3 enhances spatial learning without altering locomotion or exploratory behavior. See also figure S3. (A) Diagram of MWM with the platform in the NE quadrant during training (top) and platform removed during the probe test (bottom). (B) Time to reach the platform in seconds by day. Data represented by mean ±SEM. And analyzed via two-way ANOVA with Tukey’s multiple comparison test. *p≤0.05 (C) Time spent in each quadrant during the 60 second probe test during which the platform was removed from the NE quadrant. Data representation includes individual mice as points and bar graphs as mean ±SEM, analyzed with two-way ANOVA with Tukey’s multiple comparison test. (D) Quantification of jumping behavior observed at the conclusion of training trials. Data represented by individual mice as points and bar graphs as mean ±SEM, analyzed with unpaired T-Test. *p≤0.05 (E) Representative swim paths during probe test. In (B-E) N = 11 male *Pard3^f/f^* mice, 5 male *Pard3^f/f^:CaMKIIα-Cre* mice, 8 female *Pard3^f/f^*mice, 6 female *Pard3^f/f^:CaMKIIα-Cre* mice. (F) Diagram of the OFT chamber with outer edge vs center. (G-J) Quantification during the OFT of (G) time in outer edge, (H) number of entries into outer edge, (I) total distance traveled, and (J) average velocity. N = 6 male *Pard3^f/f^* mice, 8 male *Pard3^f/f^:CaMKIIα-Cre* mice, 13 female *Pard3^f/f^* mice, 6 female *Pard3^f/f^:CaMKIIα-Cre* mice. Data representation includes individual mice as points and bar graphs as mean ±SEM, analyzed with unpaired T-test. (K) Representative heatmap for OFT. (L) Diagram of the EPM apparatus with enclosed and open arms. (M-P) Quantification during the EPM of (M) time in open arms, (N) number of entries into open arms, (O) total distance traveled, and (P) average velocity. N = 6 male *Pard3^f/f^*mice, 8 male *Pard3^f/f^:CaMKIIα-Cre* mice, 13 female *Pard3^f/f^* mice, 6 female *Pard3^f/f^:CaMKIIα-Cre* mice. Data representation includes individual mice as points and bar graphs as mean ±SEM, analyzed with unpaired T-test. (Q) Representative heatmap for EPM.

Spatial learning and memory can be enhanced by increased anxiety-like behavior^33^. To determine if the enhanced learning and memory in *Pard3^f/f^:CaMKIIα-Cre* mice was inflated by anxiety-like behavior or increased locomotor activity, we performed the open field test (OFT) and elevated plus maze (EPM). Mice have an innate aversion to bright, open spaces. In the OFT, a square plexiglass chamber was placed under a bright light. Mice typically spend more time in the outer edge of the chamber as opposed to the open, exposed center (Figure 2F). Similarly, the EPM is a plus shaped maze consisting of two enclosed arms and two open arms elevated above the ground (Figure 2L). Usually, mice spend less time in the open, exposed arms. Both *Pard3^f/f^*and *Pard3^f/f^:CaMKIIα-Cre* entered and spent similar amounts of time in the outer edge and open arms of the OFT (Figures 2G-H and K) and EPM (Figures 2M-N and Q), respectively, indicating no aberrations in anxiety-like behavior. Similarly, there were no differences in the distance traveled or velocity of the OFT (Figures 2I-K) or EPM (Figure 2O-Q), suggesting the loss of Par3 does not alter locomotion or exploratory behavior. Together, these data demonstrate the loss of Par3 enhances spatial learning and memory without altering anxiety-like behavior, locomotion, or exploration, suggesting Par3 plays a rate-limiting role in spatial learning.

### Loss of Par3 causes dysregulation of cytoskeletal proteins and activates Rac in vivo

To explore downstream pathways of Par3, we performed unbiased phosphoproteomics of hippocampal lysates from *Pard3^f/f^* and *Pard3^f/f^:CaMKIIα-Cre* mice. Following phosphopeptide enrichment, samples were analyzed by liquid chromatography with tandem mass spectroscopy (LC-MS/MS). We identified 69 phosphosites on 67 proteins that are significantly dysregulated in the *Pard3^f/f^:CaMKIIα-Cre* as compared with littermate *Pard3^f/f^* control mice (Figure 3A and B). We then performed gene ontology (GO) analysis in Database for Annotation, Visualization, and Integrated Discovery (DAVID)^34,35^. Interestingly, we found that proteins that bind to actin or microtubules and regulate their assembly and function were among the most significantly dysregulated in the Par3 cKO hippocampus (Figure 3C-E). These data suggest that loss of Par3 leads to dysregulation of cytoskeletal dynamics in hippocampal pyramidal neurons.

**Figure 3:**
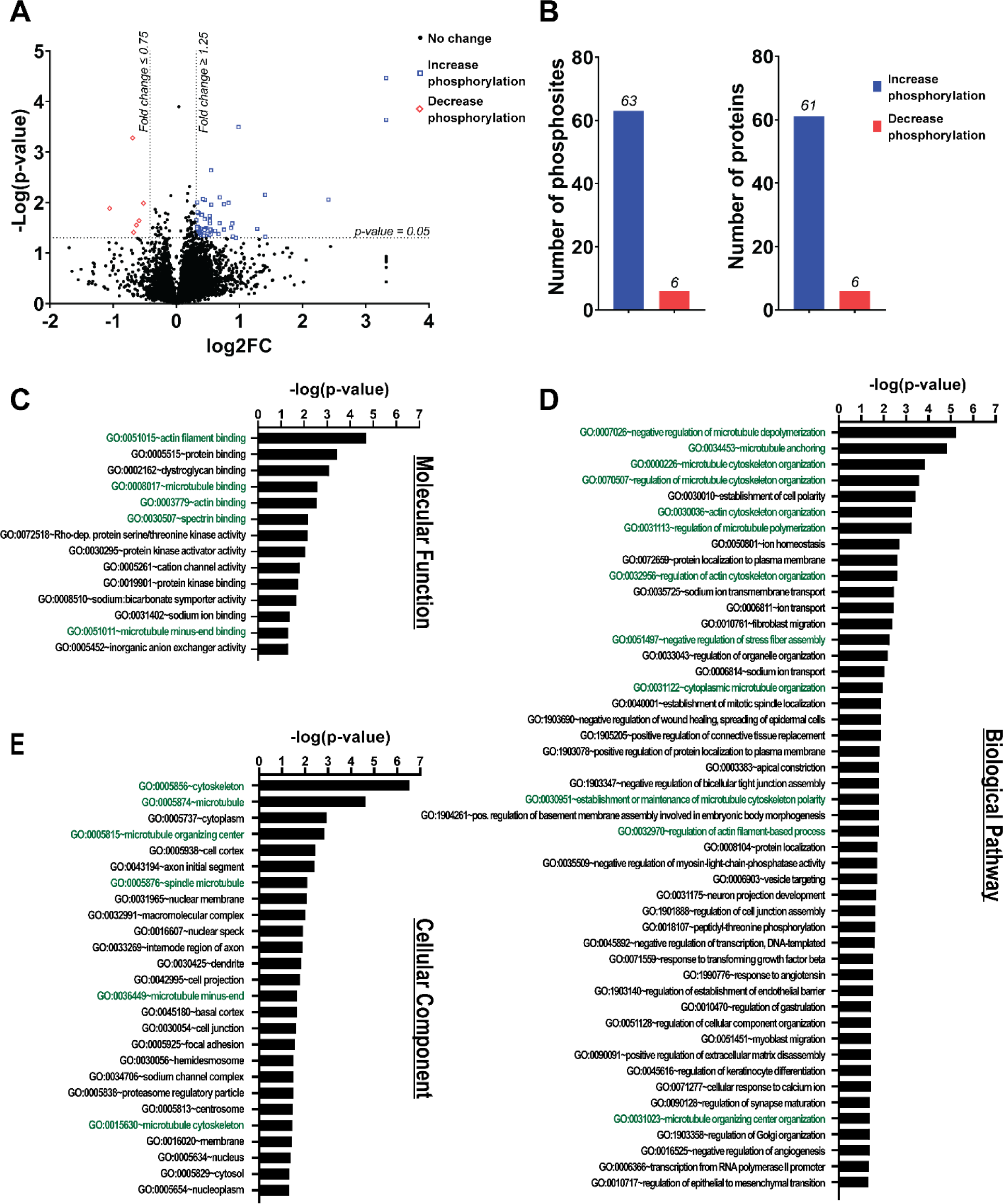
Par3 regulates phosphorylation of cytoskeleton-related proteins. (A) Phosphoproteomic analysis of hippocampal tissue. N = 3 *Pard3^f/f^* mice and 3 *Pard3^f/f^:CaMKIIα- Cre* mice, analyzed with unpaired T-test. (B) Number of phosphosites and proteins with altered phosphorylation states due to the loss of Par3. (C-D) Associated GO analysis of (C) molecular function, (D) biological pathway, and (E) cellular component of genes altered in phosphoproteomics of *Pard3^f/f^:CaMKIIα-Cre* using DAVID. Green GO terms are directly cytoskeleton related.

The observed dysregulation of cytoskeletal proteins concurs with prior studies demonstrating Par3 is upstream of TIAM1 to regulate Rac activation^13,15,36–38^, a key regulator of actin and microtubule dynamics. However, most of these studies were completed in non-neuronal cell types or in primary neuronal culture *in vitro* and report conflicting results. To investigate Rac activity *in vivo*, we used a p21-binding domain (PBD) pulldown assay to determine if Rac1 activation is altered in *Pard3^f/f^:CaMKIIα-Cre* hippocampal tissue. The PBD domain of the Rac/Cdc42 effector protein p21-activated kinase (PAK) binds GTP-bound Rac or Cdc42, enabling the detection of active Rac1 by Western blot. The loss of Par3 results in increased Rac1 activity as indicated by an ∼53% increase of active Rac1 pulled down by PAK-PBD (Figure 4A-B, p=0.0065). Furthermore, we examined the activation of PAK downstream of Rac1. We found the *Pard3^f/f^:CaMKIIα-Cre* hippocampal PSD-enriched fractions have increased phospho-PAK1 (Ser144)/PAK2 (Ser141) and no change in total PAK expression (Figure 4C-D, phospho-PAK: p=0.0077, phospho-PAK/total PAK: p=0.0074), indicating an increase in PAK activity^39^. Together, this data suggests Par3 negatively regulates Rac1 and PAK activity in the hippocampus.

**Figure 4:**
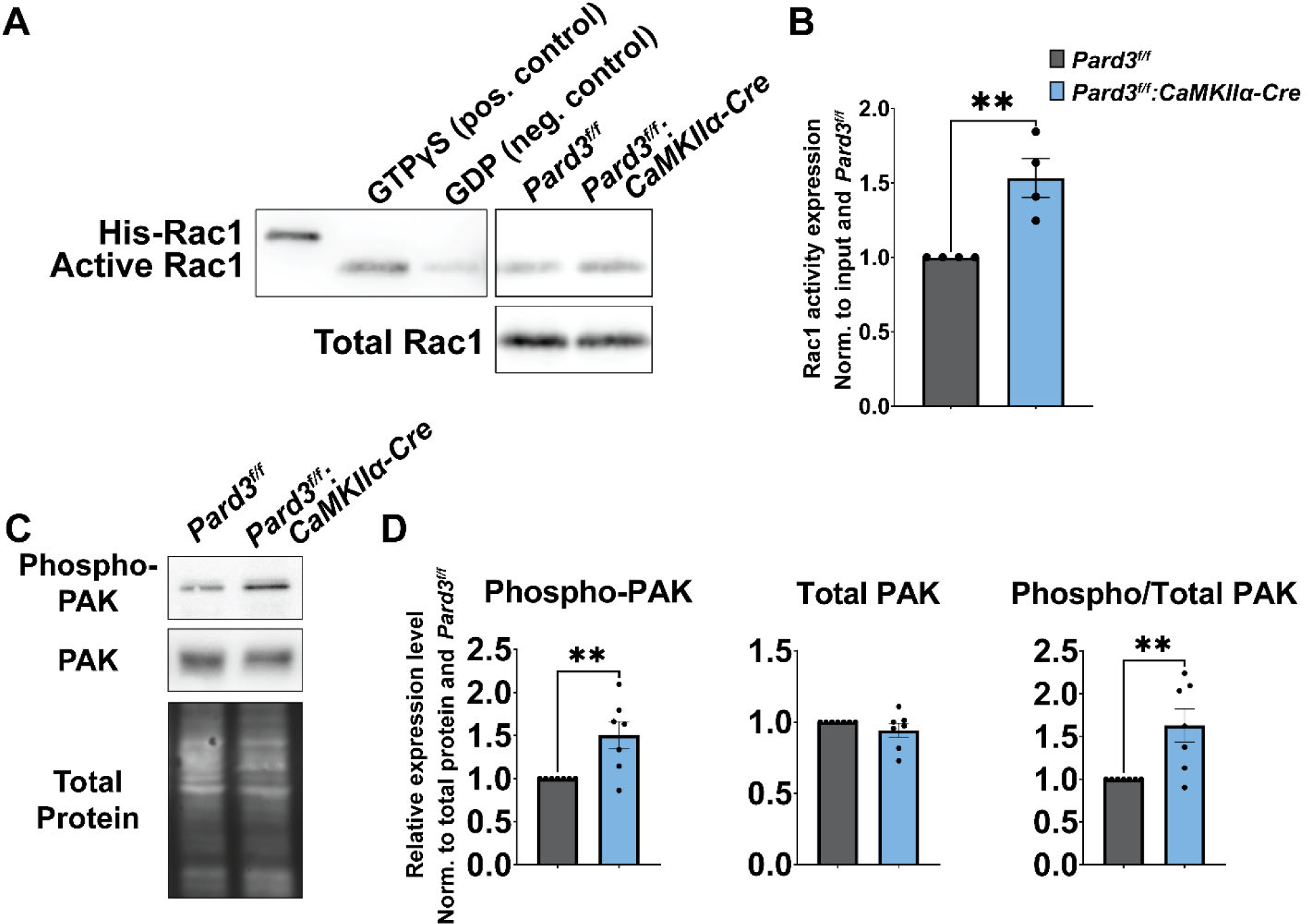
Loss of Par3 enhances the activation of the Rac1-PAK pathway. (A) Representative blots of Rac1 activity assay. (B) Quantification of Rac1 activity in hippocampal tissue normalized to total Rac1 and Pard3^f/f^. N = 4 *Pard3^f/f^* mice and 4 *Pard3^f/f^:CaMKIIα-Cre* mice. Data representation includes individual mice as points and bar graphs as mean ±SEM, analyzed with unpaired T-test. **p≤0.01 (C) Representative images for PAK-related proteins detected from PSD-enriched fractions derived from hippocampal tissue. (D) Quantification of relative expression levels normalized to total protein and *Pard3^f/f^* for phosphorylated PAK, total PAK, and phosphorylated PAK normalized to total PAK. N = 7 *Pard3^f/f^* mice and 7 *Pard3^f/f^:CaMKIIα-Cre* mice. Data representation includes individual mice as points and bar graphs as mean ±SEM, analyzed with unpaired T-test. **p≤0.01

## Discussion

In this study, we established a novel forebrain specific Par3 conditional knockout mouse model to examine the role of Par3 in dendritic spine morphogenesis and cognitive functions *in vivo*. Par3 is a key polarity regulator important for subcellular compartmentalization in several different contexts^40^. Since dendritic spines are highly compartmentalized structures^9–11,41–44^, we predicted the loss of Par3 *in vivo* would cause significant defects in the formation of mature dendritic spines, similar to the loss of mature dendritic spines in Par3 shRNA-expressing primary hippocampal neurons^15^. Unexpectedly, we observed a significant increase in dendritic spine density in the hippocampal CA1 region. The formation of mature, mushroom-shaped dendritic spines was not significantly affected. Instead, the loss of Par3 created an excess pool of long, thin dendritic spines. Consistent with the increased immature-appearing spines, we detected a significant reduction in mEPSC amplitude, suggesting a decrease in synaptic strength in the absence of Par3. These immature-appearing dendritic spines are abundant in the developing brain but are present at a lower density in adult brains. These findings suggest Par3 may serve as a molecular gatekeeper or checkpoint to limit the number of immature dendritic spines. The loss of Par3 may remove the brake on spine formation, leading to higher spine density and a larger pool of dendritic spines of immature morphology. However, it is also possible that the increase in immature dendritic spines is due to a lack of synaptic pruning, leaving an abnormally high number of immature spines.

The plasticity and stability of dendritic spines is vital to learning and memory processes^7,45,46^. Immature dendritic spines have been proposed as “learning spines”. These spines are more dynamic and can undergo potentiation during learning to encode new memories^47^. A finite number of these learning spines may be a rate-limiting step in learning and memory. Interestingly, *Pard3^f/f^:CaMKIIα-Cre* mice exhibited enhanced learning and memory in the MWM. This suggests that the increase in immature-appearing spines facilitates learning by providing a larger pool of spines capable of undergoing potentiation and thereby bypassing a rate-limiting step in learning. Notably, in patients and mouse models of neurodevelopmental disorders including Fragile X syndrome and Rett syndrome, an increase in or shift towards immature dendritic spines is often observed. This is associated with a reduction in mature dendritic spines, leading to learning impairments^48–54^. By contrast, *Pard3^f/f^:CaMKIIα-Cre* mice do not show a significant reduction in mature spines and do exhibit enhanced learning and memory. Thus, it is likely that the loss of Par3 leads to an excess pool of immature spines without impairing the transition from immature to mature spines as evident by the enhanced learning and memory abilities.

Additionally, we demonstrate that the *in vivo* depletion of Par3 leads to an elevation in the activity of the Rac1-PAK pathway. This is consistent with previous studies showing overexpression of PAK results in increased dendritic spines and synapses in primary hippocampal neurons^55^. Interestingly, a *de novo* missense mutation in *RAC1* is linked to severe intellectual disability^56^, and *PAK3* mutations are also known to be associated with intellectual disability^57–62^, indicating a key role for this pathway in higher cognition in humans. Furthermore, Rac activation has been linked to enhanced cognitive abilities in the human brain. Previous studies show the human-specific gene duplication of Slit-Robo Rho-GTPase activating protein 2 (*SRGAP2*) produces SRGAP2C, which inhibits the function of the ancestral SRGAP2A, a GTPase-activating protein (GAP) for Rac1. Thus, the presence of the human specific SRGAP2C will increase Rac1 activation. Expression of SRGAP2C in mice increases dendritic spine density, increases spine length, and enhances cognition^63,64^, which are reminiscent of the *Pard3^f/f^:CaMKIIα-Cre* mice. Considering that SNPs of Par3 have been linked to intelligence and math abilities^21,22^, it would be interesting to explore the effects of these SNPs on Par3 expression and function, and how downstream Rac1 activation may be affected.

How does Par3 regulate Rac1 activation in dendritic spines *in vivo*? Our previous studies in primary hippocampal neurons demonstrate Par3 restricts Rac1 activity to dendritic spines through interactions with the Rac guanine nucleotide exchange factor TIAM1. Loss of Par3 disperses TIAM1 and active Rac1, leading to a shift to immature dendritic spines^15^. Interestingly, TIAM1 also binds the calcium-calmodulin protein kinase II (CaMKII). After dendritic spine stimulation, CaMKII directly binds to and releases TIAM1 from an autoinhibition state, consequently activating Rac1^65,66^. In future studies, exploring the impact of Par3 regulation on CaMKII-TIAM1 interactions during synaptic plasticity may yield fascinating insights. Finally, in addition to Rac1 activation, other mechanisms may also be at play downstream of the loss of Par3. Our phosphoproteomics dataset will be an important tool for future studies aimed at understanding the signaling mechanisms of the Par polarity complex in dendritic spine plasticity as well as in other cellular processes.

In summary, we established a novel mouse model to conditionally knock out the polarity protein Par3 in forebrain pyramidal neurons. Our findings indicate that Par3 acts as a molecular checkpoint to limit immature dendritic spine density *in vivo*. The loss of Par3 leads to dendritic spine neoteny, higher spine density, and increased Rac1-PAK activity, potentially bypassing a rate-limiting step to enable enhanced learning and memory through increasing the pool of immature spines.

## Acknowledgements

The authors would like to thank Drs. David Barker, David Crockett, Bonnie Firestein, Christopher Rongo, and Loren Runnels for feedback on experimental design and manuscript. The authors would also like to thank Dr. Janet Alder for use of her MWM, Dr. Kiran Madura for the use of his microscope, and Dr. Michael Matise and Dr. Zhaohui Feng for reagents. The authors would like to thank Dr. Haiyan Zheng at the Biological Mass Spectrometry Facility at Rutgers University for completing the phosphoproteomic measurements and analysis. Cartoon diagrams were created with BioRender. This work was supported by NIH grants F31NS122477 to M.M.V., R01NS089578 to H.Z., and S10OD025140 to the Rutgers Biological Mass Spectrometry Facility.

## Author Contributions

Conceptualization, M.M.V., Z.P.P and H.Z.; Methodology, M.M.V. and M.S.; Formal Analysis, M.M.V., M.B.; Investigation, M.M.V., E.N.O., N.I., M.B. and M.S.; Writing – Original Draft, M.M.V. and H.Z.; Writing – Review & Editing, M.M.V., E.N.O., N.I., M.B., M.S., Z.P.P., H.Z.; Funding Acquisition, M.M.V. and H.Z.; Supervision, Z.P.P. and H.Z.

## Methods

### Generation and Genotyping of Pard3 conditional knockout line

To create the conditional knockout mouse line, we obtained C57BL/6N- *A^tm1Brd^Pard3^tm1a(KOMP)Wtsi^*/HMmucd (#048974-UCD) from MRC-Harwell. In this promoter driven construct line (Tm1a), a promotor driven cassette and a loxP site were inserted flanking exons 8 and 9 of *Pard3*. Exons 8 and 9 are expressed in all known variants of *Pard3*. Tm1a was crossed with a flippase line, C57BL/6N-Tg(CAG-Flpo)1Afst/Mmucd, removing the promoter driven cassette except for a loxP site, creating line Tm1c. When line Tm1c is crossed with a Cre line, Cre recombinase excises *Pard3* exons 8 and 9, rendering a nonfunctional Par3 protein. We crossed Tm1c with a CaMKIIα-Cre (Tg(Camk2a-cre)T29-1Stl; Jax#005359) to knockout Par3 in postnatal forebrain pyramidal neurons during synaptogenesis. Genotypes were determined via PCR of DNA collected from ear or tail tissue. For the presence of floxed *Pard3*, the following primers were used: 5arm-WTF (5’-CTGTTATCCTCCAAACCCTGA-3’), Crit-WTR (5’- ACACTGGGAGAGACCACCAC-3’), and 5arm-MutR (5’-GAACTTCGGAATAGGAACTTCG-3’).

For the presence of *CaMKIIα-Cre*, the following primers were used: Forward (5’- GAACCTGATGGACATGTTCAGG-3’) and Reverse (5’-AGTGCGTTCGAACGCTAGAGCCTGT-3’). Cre expression was validated by crossing the *Pard3* line with B6.129X1- *Gt(ROSA)26Sor^tm1(EYFP)Cos^*/J. Mice were housed in a 12 hour dark/light cycle with ad libitum access to chow and water. Experiments were completed in male and female 5- to 6-week-old mice, unless otherwise indicated. All experiments are in compliance with the Rutgers University institutional animal care and use committee (IACUC) protocols.

### Golgi Staining, imaging, and analysis

Brains were processed using FD Rapid GolgiStain kit (FD NeuroTechnologies, Inc., PK401) per manufacturer’s directions. Briefly, mice were deeply anesthetized with 1.2% Avertin. After non-response to tail and toe pinch, mice were decapitated. The brain was removed and placed in Solution A + B mixed 24 hours prior. After an overnight incubation at room temperature in the dark, the Solution A + B was changed. After a 3-week total incubation in the dark, the brain was transferred to Solution C for 72 hours with a change after 24 hours. Brains were rapidly frozen in pre-chilled isopentane for 5 minutes on dry ice and stored at -80°C until sectioning. 200µm sections were cut using a Cryostat (Microm HM505E) and plated on gelatin coated slides. After drying overnight in the dark the tissue was rinsed twice in ddH_2_O, stained with diluted Solution D + E, rinsed twice in ddH_2_O, dehydrated in 50%, 75%, 95% ethanol, further dehydrated four times in 100% ethanol, cleared thrice in xylene, and coverslipped with Permount (Fisher Scientific, SP15-100). Z-stack images were collected at 0.25 µm thickness using either a Leica DMRE using the Hamamatsu ORCA-ER camera and Leica 506185 63x objective or Axio Imager M1 Zeiss using the ZxioCam MRM camera, AxioVision Rel. 4.8 program, and 100X Plan-APOCHROMAT 100X/1,40:|DIC infinity/0.17 objective. Images were processed and dendritic spines analyzed in FIJI^67^ and RECONSTRUCT^68,69^. Morphology categorical cutoffs were used as previously described^69^. Outliers were calculated and removed according to ROUT with Q value of 1%.

### Behavior

All mice were handled for 3 days prior to the start of behavioral tests. A handling tunnel was used to minimize stress^70^. Mice were moved to the testing room at least 30 minutes prior to experimentation on all behavioral test days. MWM was analyzed by EthoVision XT. EPM and OFT were analyzed by AnyMaze.

### Morris Water Maze (MWM)

The MWM pool is 1.3 meter in diameter filled with opaque water maintained at 21±1°C. During testing the cages were placed on a warming pad. During the 4 training days mice received 5 trials per day with at least 15-minute intertrial intervals. The trial starting location pattern was never repeated across the 4 training days. Mice were released into the water facing the pool wall and were guided to the platform if they do not reach it within 60 seconds. Mice remained on the platform for 15 seconds before being retrieved. On day 5, the probe test is completed by removing the platform, releasing the mice from a novel location, and allowing the mice to swim for 60 seconds. Latency to platform, time in quadrant, velocity, and cumulative distance from platform were measured using EthoVision XT.

### Elevated Plus Maze (EPM)

The EPM apparatus was 56 cm above the ground. The arms were 5 cm wide and 25 cm long. The enclosed arms had 16 cm high walls. Mice were placed in the center of the apparatus and allowed to freely explore for 10 minutes.

### Open Field Test (OFT)

The OFT was conducted in a 40 cm x 40 cm x 40 cm white plexiglass box with a house light positioned directly above the apparatus. Mice were placed in the center of the apparatus and allowed to freely explore for 10 minutes.

### Primary Neuronal Culture

Cultures were harvested and maintained as previously described^71^. Briefly, *Pard3^f/f^* males and females were bred, and neurons were harvested from E17.5 embryonic brains. High density cultures were seeded at 8.5x10^5^ cells per 35 mm dish coated with 0.1 mg/mL poly-L-lysine. At DIV0 cells were infected with either AAV-Cre-GFP-hSyn (Addgene, 105540-AAV9) or AAV-GFP (Addgene, 105539-AAV9). At DIV14, neurons were lysed in RIPA buffer (20mM Tris-HCl pH 7.4, 150mM NaCl, 0.5% NP-40, 1.0% Triton-X-100, 2 mM EDTA, 2mM EGTA, 0.25% NaDOC).

Lysates were cleared at 16,100 x g for 10 minutes at 4°C and analyzed via Western Blot.

### Synaptosomal fractionation, Western blots

For all synaptosomal fractions and Rac activity assays, mice were euthanized via cervical dislocation and the hippocampi were dissected in ice-cold 1xPBS and flash frozen on dry ice. For synaptosomal fractionation, tissue was homogenized in 10mM HEPES pH 7.4, 2mM EDTA, protease inhibitor (Sigma, P-8340), and phosphatase inhibitor (Sigma, P0044) using a 2 mL glass douncer and incubated on ice for 10 minutes. Lysates were centrifuged for 10 minutes at 1,000 x g and 4°C creating the P1 nuclei pellet and S1 homogenate supernatant. The S1 homogenate supernatant was separated and centrifuged for 15 minutes at 10,000 x g and 4°C creating the S2 cytosolic supernatant and the P2 crude synaptosome pellet. S2 was removed. P2 was resuspended in 50mM HEPES pH 7.4, 2mM EDTA, 2mM EGTA, 1% Triton-X-100, protease inhibitor (Sigma, P-8340), and phosphatase inhibitor (Sigma, P0044). Resuspended P2 was centrifuged for 80 minutes at 20,000 x g and 4°C creating the S3 vesicular/presynaptic supernatant and the P3 PSD-enriched pellet. P3 was resuspended in 100mM Tris pH 9, 1% sodium deoxycholate, 0.025% SDS, protease inhibitor cocktail (Sigma, P-8340), and phosphatase inhibitor cocktail (Sigma, P0044). Fraction concentrations were measured with Pierce 600nm Protein Assay Reagent (Thermo Scientific, 22660) and prepared in Laemmli’s sample buffer.

Rac1 activity was determined using the Rac1 Pull-Down Activation Assay Biochem Kit (Cytoskeleton, Inc., BK035), following manufacture’s protocol. Briefly, 300 µg of hippocampal lysate was incubated with 10 µg of GST-tagged PAK-PBD beads, washed, and analyzed via Western Blot. For positive and negative controls 300 µg of protein was incubated with either GTPγS or GDP, respectively, for 15 minutes at room temperature immediately prior to incubation with GST-PAK-PBD beads.

Protein expression levels were analyzed via Western Blot analysis. Synaptosomal protein expression levels were normalized to total protein using No-Stain Protein Labeling Reagent (Invitrogen, A44449). Primary antibodies (Anti-Shank3 (N367/62), 1:500, NeuroMAB 75-344; Anti-Homer1, 1:1,000, Synaptic Systems, 160 003; Anti-GluR1 (N355/1), 1:2,000, UC Davis,73,327; Anti-GluR2, 1:4,000, NeuroMAB, MAB397; Anti-PSD-95, 1:2,000, UC Davis, K28/43; Anti-NMDAR1, 1:1,000, BD Pharmingen, 5563008; Anti-Rac1, 1:500, Cytoskeleton, Inc., ARC03; Anti-Phospho PAK, 1:2,000, Cell Signaling Technologies, 2606; Anti-PAK, 1:2,000, Santa Cruz, 166887) were diluted in 1 x PBS + 0.15% (v/v) Tween-20 and 3% (w/v) bovine serum albumin. Secondary antibodies (Peroxidase-conjugated AffiniPure Goat Anti-Rabbit IgG, 1:5,000, Jackson ImmunoResearch Laboratories, Inc., 111-035-003; Peroxidase-conjugated AffiniPure Goat Anti-Mouse IgG, 1:5,000, Jackson ImmunoResearch Laboratories, Inc., 115-035-003) were diluted in 1 x PBS + 0.15% Tween-20 and 3% (w/v) bovine serum albumin. Densitometry was measured using FIJI^67^.

### Electrophysiology

Brain slice physiology was performed in the dorsal CA1 pyramidal neurons in 5-7-week-old mice as described previously with some modifications (Pang et al. 2002). Mice were anesthetized and euthanized by intracardial perfusion with ice-cold cutting solution (125mM NaCl, 25mM NaHCO_3_, 2.5mM KCl, 1.25mM NaH_2_PO_4_, 2mM CaCl_2_, 10mM MgCl_2_, 2.5mM glucose) oxygenated with 95% O_2_/5% CO_2_. 300 µm sagittal sections were cut in ice-cold cutting solution oxygenated with 95% O_2_/5% CO_2_. Slices were transferred to cutting solution at 33°C for 10 minutes and then to a second warm beaker filled with ACSF (125mM NaCl, 25mM NaHCO_3_, 2.5mM KCl, 1.25mM NaH_2_PO_4_, 2.5mM CaCl_2_, 1.2mM MgCl_2_, and 2.5mM glucose) oxygenated with 95% O_2_/5% CO_2_. Post 1 hour recovery, sections were perfused continuously at 4 mL/minute with oxygenated ACSF plus 22.5mM sucrose in a recording chamber at room temperature (∼30°C). During recordings, a CA1 neuron was held at -70mV and recorded in voltage-clamp mode for 3-5 minutes while in ACSF plus 50µM picrotoxin (PTX), 40µM AP5, and 1µM tetrodotoxin (TTX). Traces were analyzed in Clampfit (Molecular Devices) and filtered using a lowpass boxcar filter with 11 smoothing points with a manual template search for mEPSCs. Events below 10 pA were excluded. Amplitudes were determined by averaging all events in a single recording.

### Phosphoproteomic analysis

Hippocampal tissue was lysed in 8M Guanidine HCl, 50mM Tris, pH7.5, 5mM DTT and processed for phosphoproteomic analysis as previously described^72^. Database for Annotation, Visualization, and Integrated Discovery (DAVID)^34,35^ was used to analyze molecules based on Functional GO Analysis.

### Statistical analysis

For manual measurements samples were blinded at both collection and analysis. Multiple data points collected within the same animal (dendritic spine density and morphology data) were analyzed in Statistical Analysis System (SAS) using a mixed model with simple covariance to nest the multiple measurements within subjects as previously described^73^. All other data sets were analyzed in Prism GraphPad using an unpaired T-test or two-way ANOVA with Tukey’s multiple comparison tests. Graphs were made in GraphPad Prism. Bar graphs depict individual measurements as data points and mean ±standard error of the mean (SEM). Violin plots depict thick lines as median and dotted lines as quartiles. Sample size details are listed in figure legends. Statistical significance was set at p ≤ 0.05.

## Supplemental Figures and Legends

Figure S1: Generation of *Pard3* conditional knockout mouse model

(A) Schematic of strategies in creating conditional allele of *Pard3* knockout mouse line. Tm1a line crossed with flippase creates Tm1c line. Tm1c line crossed with *CaMKIIa-Cre* line creates Tm1d line knocking out exons 8 and 9 of *Pard3*.

(B) DNA gel of genotyping results for *Pard3^+/+^, Pard3^f/+^,* and *Pard3^f/f^*.

(C) *Pard3^f/f^*and *Pard3^f/f^:CaMKIIα-Cre* mice crossed with B6.129X1-Gt(ROSA)26Sor^tm1(EYFP)Cos^/J (R26R-EYFP) express enhanced yellow fluorescent protein in Cre-expressing cells (green) and stained with DAPI (blue) in the (upper) medial prefrontal cortex (mPFC), (middle) CA1, and (lower) dentate gyrus (DG). Scale bar = 100µm.

(D) Par3 is knocked out using AAV-hSyn-Cre applied to primary neuronal culture derived from *Pard3^f/f^* E17.5 hippocampi.

(E) Body weight of 5-week-old mice. N = 12 male *Pard3^f/f^*, 11 male *Pard3^f/f^:CaMKIIα-Cre*, 8 female *Pard3^f/f^,* 11 female *Pard3^f/f^:CaMKIIα-Cre*. Data representation includes individual mice as points and bar graphs as mean ±SEM, analyzed with two-way ANOVA with Tukey’s multiple comparisons test.

Figure S2 related to figure 1: Validation of synaptosomal investigation.

(A) Image of hippocampus of Golgi-stained tissue. Scale bar = 210µm.

(B) Image of CA1 pyramidal neuron. Scale bar = 25µm.

(C) Schematic of tissue fractionation to generate PSD-enriched fractions.

(D) Western blot images of all fractionation samples of *Pard3^f/f^* hippocampal tissue including postsynaptic density protein 95 (PSD-95), synapsin, synaptic vesicle protein 2 (SV2), glyceraldehyde-3-phosphate dehydrogenase (GAPDH), Glial fibrillary acidic protein (GFAP), Ionized calcium binding adaptor molecule 1 (IBA1), and lysosomal associated membrane protein- 2 (LAMP2).

Figure S3 related to Figure 2: Additional characterization of MWM performance in *Pard3^f/f^* and *Pard3^f/f^:CaMKIIα-Cre* Mice.

(A-B) Comparison of males vs females in latency to platform for (A) *Pard3^f/f^* and (B) *Pard3^f/f^:CaMKIIα-Cre*.

(C) Average velocity of swimming speed for each day of the MWM.

(D) Time in each quadrant during the probe test of the MWM comparing each quadrant within *Pard3^f/f^* and *Pard3^f/f^:CaMKIIα-Cre*.

(E) Cumulative distance from the platform during the probe test. In (A-E) N = 14 male *Pard3^f/f^*, 5 male *Pard3^f/f^:CaMKIIα-Cre*, 12 female *Pard3^f/f^*, 6 female *Pard3^f/f^:CaMKIIα-Cre*. (A-D) Analyzed via two-way ANOVA with Tukey’s multiple comparison test.

(E) Analyzed by Student’s T-Test. ***p<0.001, ****p < 0.0001

## References

1. C H Bailey, a. & Kandel, E.R. Structural Changes Accompanying Memory Storage. Annual review of physiology 55, 397–426 (1993).

2. Huang, L., Zhou, H., Chen, K., Chen, X. & Yang, G. Learning-Dependent Dendritic Spine Plasticity Is Reduced in the Aged Mouse Cortex. Front Neural Circuits 14, 581435 (2020).

3. Hofer, S.B., Mrsic-Flogel, T.D., Bonhoeffer, T. & Hübener, M. Experience leaves a lasting structural trace in cortical circuits. Nature 457, 313–317 (2009).

4. Xu, T., et al. Rapid formation and selective stabilization of synapses for enduring motor memories. Nature 462, 915–919 (2009).

5. Kim, S.K. & Nabekura, J. Rapid synaptic remodeling in the adult somatosensory cortex following peripheral nerve injury and its association with neuropathic pain. The Journal of neuroscience : the official journal of the Society for Neuroscience 31, 5477–5482 (2011).

6. Lai, C.S., Franke, T.F. & Gan, W.B. Opposite effects of fear conditioning and extinction on dendritic spine remodelling. Nature 483, 87–91 (2012).

7. Hayashi-Takagi, A., et al. Labelling and optical erasure of synaptic memory traces in the motor cortex. Nature 525, 333–338 (2015).

8. Yang, G., Pan, F. & Gan, W.-B. Stably maintained dendritic spines are associated with lifelong memories. Nature 462, 920–924 (2009).

9. Colgan, L.A. & Yasuda, R. Plasticity of dendritic spines: subcompartmentalization of signaling. Annual review of physiology 76, 365–385 (2014).

10. Hotulainen, P. & Hoogenraad, C.C. Actin in dendritic spines: connecting dynamics to function. J Cell Biol 189, 619–629 (2010).

11. Spence, E.F. & Soderling, S.H. Actin Out: Regulation of the Synaptic Cytoskeleton. J Biol Chem 290, 28613–28622 (2015).

12. Rodriguez-Boulan, E. & Macara, I.G. Organization and execution of the epithelial polarity programme. Nat Rev Mol Cell Biol 15, 225–242 (2014).

13. Landin Malt, A., et al. Par3 is essential for the establishment of planar cell polarity of inner ear hair cells. Proc Natl Acad Sci U S A 116, 4999–5008 (2019).

14. Chuykin, I., Ossipova, O. & Sokol, S.Y. Par3 interacts with Prickle3 to generate apical PCP complexes in the vertebrate neural plate. Elife 7(2018).

15. Zhang, H. & Macara, I.G. The polarity protein PAR-3 and TIAM1 cooperate in dendritic spine morphogenesis. Nat Cell Biol 8, 227–237 (2006).

16. Zhang, H. & Macara, I.G. The PAR-6 polarity protein regulates dendritic spine morphogenesis through p190 RhoGAP and the Rho GTPase. Dev Cell 14, 216–226 (2008).

17. Duman, J.G., et al. The adhesion-GPCR BAI1 regulates synaptogenesis by controlling the recruitment of the Par3/Tiam1 polarity complex to synaptic sites. The Journal of neuroscience : the official journal of the Society for Neuroscience 33, 6964–6978 (2013).

18. Scott, J., et al. Apical-Basal Polarity Signaling Components, Lgl1 and aPKCs, Control Glutamatergic Synapse Number and Function. iScience 20, 25-41 (2019).

19. Pinto, D., et al. Convergence of Genes and Cellular Pathways Dysregulated in Autism Spectrum Disorders. The American Journal of Human Genetics 94, 677–694 (2014).

20. Kim, S.K., Lee, J.Y., Park, H.J., Kim, J.W. & Chung, J.-H. Association study between polymorphisms of the PARD3 gene and schizophrenia. Exp Ther Med 3, 881–885 (2012).

21. Davies, G., et al. Study of 300,486 individuals identifies 148 independent genetic loci influencing general cognitive function. Nature Communications 9, 2098 (2018).

22. Lee, J.J., et al. Gene discovery and polygenic prediction from a genome-wide association study of educational attainment in 1.1 million individuals. Nature Genetics 50, 1112–1121 (2018).

23. Wang, T., et al. De novo genic mutations among a Chinese autism spectrum disorder cohort. Nat Commun 7, 13316 (2016).

24. Stessman, H.A., et al. Targeted sequencing identifies 91 neurodevelopmental-disorder risk genes with autism and developmental-disability biases. Nat Genet 49, 515–526 (2017).

25. Guo, H., et al. Inherited and multiple de novo mutations in autism/developmental delay risk genes suggest a multifactorial model. Mol Autism 9, 64 (2018).

26. Zhou, X., et al. Integrating de novo and inherited variants in 42,607 autism cases identifies mutations in new moderate-risk genes. Nat Genet 54, 1305–1319 (2022).

27. Anney, R., et al. Individual common variants exert weak effects on the risk for autism spectrum disorders. Human molecular genetics 21, 4781–4792 (2012).

28. Insolera, R., Chen, S. & Shi, S.H. Par proteins and neuronal polarity. Dev Neurobiol 71, 483–494 (2011).

29. Tsien, J.Z., et al. Subregion- and cell type-restricted gene knockout in mouse brain. Cell 87, 1317–1326 (1996).

30. Kim, I.H., Wang, H., Soderling, S.H. & Yasuda, R. Loss of Cdc42 leads to defects in synaptic plasticity and remote memory recall. Elife 3(2014).

31. Shors, T.J., Chua, C. & Falduto, J. Sex differences and opposite effects of stress on dendritic spine density in the male versus female hippocampus. The Journal of neuroscience : the official journal of the Society for Neuroscience 21, 6292–6297 (2001).

32. Won, H., et al. Autistic-like social behaviour in Shank2-mutant mice improved by restoring NMDA receptor function. Nature 486, 261–265 (2012).

33. Eysenck, M.W. Anxiety, learning, and memory: A reconceptualization. Journal of Research in Personality 13, 363–385 (1979).

34. Huang da, W., Sherman, B.T. & Lempicki, R.A. Systematic and integrative analysis of large gene lists using DAVID bioinformatics resources. Nat Protoc 4, 44–57 (2009).

35. Huang da, W., Sherman, B.T. & Lempicki, R.A. Bioinformatics enrichment tools: paths toward the comprehensive functional analysis of large gene lists. Nucleic Acids Res 37, 1–13 (2009).

36. Chen, X. & Macara, I.G. Par-3 controls tight junction assembly through the Rac exchange factor Tiam1. Nat Cell Biol 7, 262–269 (2005).

37. Nishimura, T., et al. PAR-6-PAR-3 mediates Cdc42-induced Rac activation through the Rac GEFs STEF/Tiam1. Nat Cell Biol 7, 270–277 (2005).

38. Duman, J.G., et al. The adhesion-GPCR BAI1 regulates synaptogenesis by controlling the recruitment of the Par3/Tiam1 polarity complex to synaptic sites. J Neurosci 33, 6964–6978 (2013).

39. Chong, C., Tan, L., Lim, L. & Manser, E. The mechanism of PAK activation. Autophosphorylation events in both regulatory and kinase domains control activity. J Biol Chem 276, 17347–17353 (2001).

40. Goldstein, B. & Macara, I.G. The PAR proteins: fundamental players in animal cell polarization. Developmental cell 13, 609–622 (2007).

41. Bloodgood, B.L. & Sabatini, B.L. Neuronal activity regulates diffusion across the neck of dendritic spines. *Science (New York*, N.Y.) 310, 866–869 (2005).

42. Gulledge, A.T., Carnevale, N.T. & Stuart, G.J. Electrical advantages of dendritic spines.PLoS One 7, e36007 (2012).

43. Svoboda, K., Tank, D.W. & Denk, W. Direct measurement of coupling between dendritic spines and shafts. *Science (New York*, N.Y.) 272, 716–719 (1996).

44. Yuste, R. Electrical compartmentalization in dendritic spines. Annual review of neuroscience 36, 429–449 (2013).

45. Grutzendler, J., Kasthuri, N. & Gan, W.B. Long-term dendritic spine stability in the adult cortex. Nature 420, 812–816 (2002).

46. Trachtenberg, J.T., et al. Long-term in vivo imaging of experience-dependent synaptic plasticity in adult cortex. Nature 420, 788–794 (2002).

47. Bourne, J. & Harris, K.M. Do thin spines learn to be mushroom spines that remember? Current opinion in neurobiology 17, 381–386 (2007).

48. Hinton, V.J., Brown, W.T., Wisniewski, K. & Rudelli, R.D. Analysis of neocortex in three males with the fragile X syndrome. Am J Med Genet 41, 289–294 (1991).

49. Wisniewski, K.E., Segan, S.M., Miezejeski, C.M., Sersen, E.A. & Rudelli, R.D. The Fra(X) syndrome: neurological, electrophysiological, and neuropathological abnormalities. Am J Med Genet 38, 476–480 (1991).

50. Rudelli, R.D., et al. Adult fragile X syndrome. Clinico-neuropathologic findings. Acta neuropathologica 67, 289–295 (1985).

51. He, C.X. & Portera-Cailliau, C. The trouble with spines in fragile X syndrome: density, maturity and plasticity. Neuroscience 251, 120–128 (2013).

52. Belichenko, P.V., et al. Widespread changes in dendritic and axonal morphology in Mecp2-mutant mouse models of Rett syndrome: evidence for disruption of neuronal networks. The Journal of comparative neurology 514, 240–258 (2009).

53. Armstrong, D.D. The neuropathology of the Rett syndrome. Brain Dev 14 **Suppl**, S89-98 (1992).

54. Nakai, N., Takumi, T., Nakai, J. & Sato, M. Common Defects of Spine Dynamics and Circuit Function in Neurodevelopmental Disorders: A Systematic Review of Findings From in Vivo Optical Imaging of Mouse Models. Front Neurosci 12, 412 (2018).

55. Zhang, H., Webb, D.J., Asmussen, H., Niu, S. & Horwitz, A.F. A GIT1/PIX/Rac/PAK Signaling Module Regulates Spine Morphogenesis and Synapse Formation through MLC. The Journal of Neuroscience 25, 3379–3388 (2005).

56. Lelieveld, S.H., et al. Meta-analysis of 2,104 trios provides support for 10 new genes for intellectual disability. Nat Neurosci 19, 1194–1196 (2016).

57. Meng, J., Meng, Y., Hanna, A., Janus, C. & Jia, Z. Abnormal Long-Lasting Synaptic Plasticity and Cognition in Mice Lacking the Mental Retardation Gene *Pak3*. The Journal of Neuroscience 25, 6641–6650 (2005).

58. Bienvenu, T., et al. Missense mutation in PAK3, R67C, causes X-linked nonspecific mental retardation. Am J Med Genet 93, 294–298 (2000).

59. Allen, K.M., et al. PAK3 mutation in nonsyndromic X-linked mental retardation. Nat Genet 20, 25–30 (1998).

60. Hayashi, M.L., et al. Inhibition of p21-activated kinase rescues symptoms of fragile X syndrome in mice. Proc Natl Acad Sci U S A 104, 11489–11494 (2007).

61. Nagy, D., et al. Further delineation of the phenotype of PAK3-associated x-linked intellectual disability: Identification of a novel missense mutation and review of literature. Eur J Med Genet 63, 103800 (2020).

62. Iida, A., et al. A novel PAK3 pathogenic variant identified in two siblings from a Japanese family with X-linked intellectual disability: case report and review of the literature. Cold Spring Harb Mol Case Stud 5(2019).

63. Fossati, M., et al. SRGAP2 and Its Human-Specific Paralog Co-Regulate the Development of Excitatory and Inhibitory Synapses. Neuron 91, 356–369 (2016).

64. Schmidt, E.R.E., Kupferman, J.V., Stackmann, M. & Polleux, F. The human-specific paralogs SRGAP2B and SRGAP2C differentially modulate SRGAP2A-dependent synaptic development. Sci Rep 9, 18692 (2019).

65. Kojima, H., et al. The role of CaMKII-Tiam1 complex on learning and memory. Neurobiology of Learning and Memory 166, 107070 (2019).

66. Saneyoshi, T., et al. Reciprocal Activation within a Kinase-Effector Complex Underlying Persistence of Structural LTP. Neuron 102, 1199–1210.e1196 (2019).

67. Schindelin, J., et al. Fiji: an open-source platform for biological-image analysis. Nature Methods 9, 676–682 (2012).

68. Fiala, J.C. Reconstruct: a free editor for serial section microscopy. J Microsc 218, 52–61 (2005).

69. Risher, W.C., Ustunkaya, T., Singh Alvarado, J. & Eroglu, C. Rapid Golgi analysis method for efficient and unbiased classification of dendritic spines. PLoS One 9, e107591 (2014).

70. Gouveia, K. & Hurst, J.L. Optimising reliability of mouse performance in behavioural testing: the major role of non-aversive handling. Scientific Reports 7, 44999 (2017).

71. Sun, M., Bernard, L.P., Dibona, V.L., Wu, Q. & Zhang, H. Calcium phosphate transfection of primary hippocampal neurons. Journal of visualized experiments : JoVE, e50808 (2013).

72. Retzbach, E.P., et al. Independent effects of Src kinase and podoplanin on anchorage independent cell growth and migration. Mol Carcinog 61, 677–689 (2022).

73. Wilson, M.D., Sethi, S., Lein, P.J. & Keil, K.P. Valid statistical approaches for analyzing sholl data: Mixed effects versus simple linear models. Journal of neuroscience methods 279, 33–43 (2017).

